# A Patient-Specific Model of Transcatheter Valve Replacement in a Bicuspid Heart Valve

**DOI:** 10.1101/2021.03.05.433633

**Authors:** Orla M. McGee, Adrian McNamara, Laoise M. McNamara

**Affiliations:** Biomechanics Research Centre, Biomedical Engineering, College of Engineering & Informatics, National University of Ireland Galway; Boston Scientific, Ballybrit Business Park, Ballybrit, Galway

## Abstract

Bicuspid Aortic Valves (BAVs) are a common congenital heart disease where two cusps of the aortic heart valve become fused together, this leads to two unequally sized leaflets compared to the normal trileaflet valve. Transcatheter Aortic Valves are currently used in off-label treatmet of stenosed BAVs, however, due to the abnormal valve anatomy, debate surrounds the sizing of transcatheter valves. In this study, finite element models were developed to simulate the deployment of two different valves sizes (a 25 mm and a 27 mm) of the Lotus valve into the patient-specific aortic root geometry of a clinical stenosed BAV case. These models were used to investigate and compare the eccentricity, stress and mal-apposition of the two valve sizes. The results demonstrated that the 25 mm valve was the most suitable in terms of eccentricity and stress reduction. It was also shown that the smaller 25 mm valve size did not increase the likelihood of mal-apposition. As the 25 mm valve was deemed suitable based on current sizing algorithms, on the basis of these results traditional annulus measurement and device sizing may be suitable in the case of the Lotus valve.

## Introduction

Bicuspid Aortic Valves (BAVs) are a common congenital cardiac malformation of the aortic valve, where two cusps of the valve have become fused together due to a rheumatic or inflammatory process [1, 2]. BAVs are the most common congenital valve abnormality and have been found to occur in 0.5-2% of patients [3–6]. Studies have suggested that >20% of patients requiring TAVI procedures are BAV patients [7, 8]. A BAV is comprised of two, often unequally sized, leaflets instead of the normal tri-leaflet aortic valve. Figure 1 shows a schematic of the different anatomic classifications of BAVs. It has been estimated that 90% of BAVs are Type I where the left and right leaflets are fused together [9].

**Figure 1:**
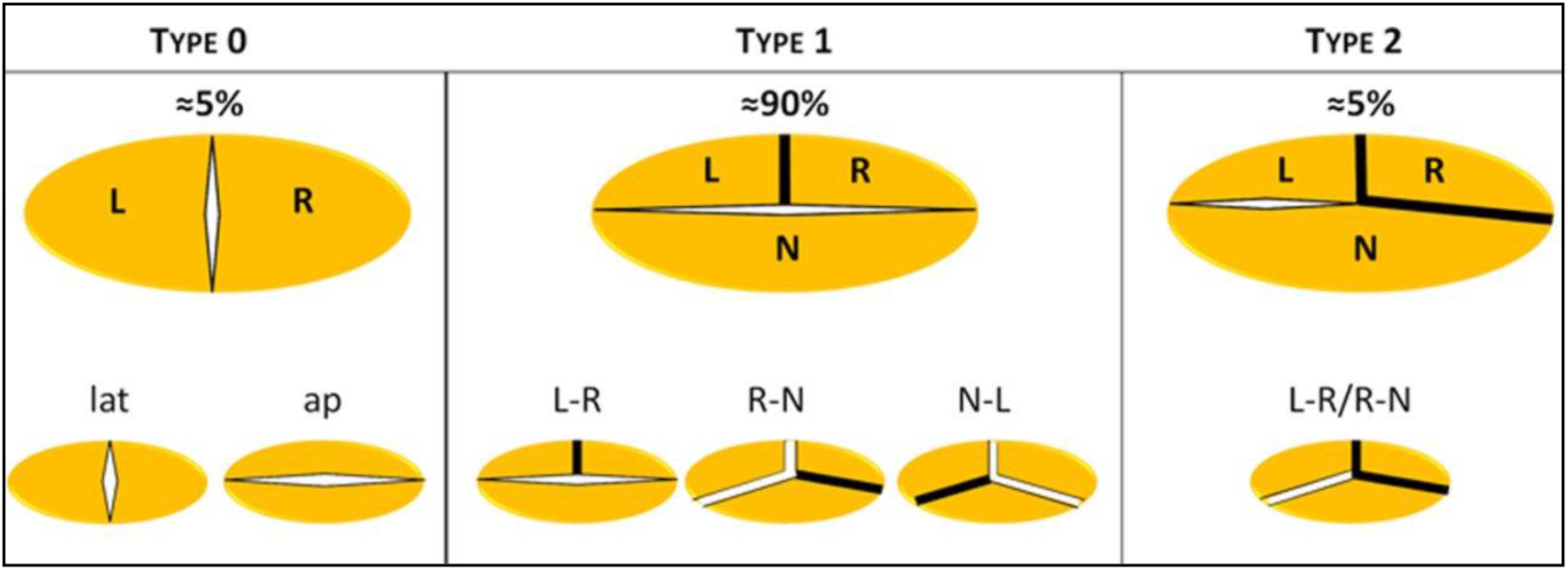
Schematic of the anatomical classification of BAVs where the prominent black line represents the fusion of the leaflets L, left coronary sinus; lat, lateral; N, noncoronary sinus; and R, right coronary sinus [9].

The clinical trials that established TAVI as the standard treatment in inoperable patients excluded BAV patients [10–14]. Further to this, the treatment of BAV was contraindicated for earlier generation valves due to concerns regarding the elliptical anatomy of BAVs leading to valve malfunction and valve positioning [15–17]. This is due to the abnormal valve geometry of BAVs, which are commonly associated with eccentricity and factors such as asymmetrical valve calcification, difference in leaflet sizes and concomitant aortopathy [18–22]. In BAV patients treated with TAV, reports have associated BAVs with malfunction, mal-apposition, incomplete sealing, severe PVL and aortic regurgitation [16, 17, 23–25]. However, due to increased experience and advances in technology, off-label treatment of BAV patients using TAVI is increasing [26]. Despite the higher rates of PVL and lower device success in earlier generation valves, new generation valves have shown improved outcomes in the treatment of BAV patients with less PVL [27, 28]. It was reported that there was no significant difference between BAV patients and tricuspid aortic valve patients in terms of 30-day mortality, mean peak gradients, PVL, need for pacemaker implantation or life-threatening bleeding [9, 15]. However, it has been argued that none of the reported BAV cohorts have adequate sample sizes, motivating the need for further studies [22]. Furthermore, appropriate sizing of TAVI for treatment of BAV patients has not been determined. PVL has been particularly problematic in relation to BAV patients, a common complication which has been shown to reduce with oversizing [29–32]. It has previously been shown that using the SAPIEN 3 over the SAPIEN XT reduces the need for oversizing due to the sealing skirt. However, high percentages (2.3%) of annulus rupture occurred using the SAPIEN 3. Recent expert analysis has proposed that traditional (i.e. sizing developed for tricuspid aortic valve patients) annulus measurement and device sizing may not necessarily be appropriate in treating BAV stenosis [10]. Furthermore, the implications of underexpansion of valves in BAV cases in terms of structural failure of the valve are unknown due to limited data existing on long-term durability in BAV cases [10]. Moreover, BAV studies have not used multi-detector CT post-TAVI, limiting the opportunity to assess and learn from post-deployment configuration of the bioprosthetic devices [9]. Due to limited proof of efficacy, additional research is needed to investigate the efficacy of TAVI in the treatment of BAV stenosis [22, 26]. As TAVI progresses to treat lower-risk patients, the treatment of BAV stenosis using TAVI is becoming one of the most topical yet promising ventures in the field of cardiovascular intervention. This is heightened by the fact that BAVs are more prevalent in younger patient cohorts [22, 33] and bicuspid aortic stenosis typically presents itself a decade or earlier than that of tri-leaflet patients [34].

The Lotus valve is a next generation repositionable valve with a unique mechanical locking mechanism that shortens longitudinally and expands radially during deployment. It does not require rapid pacing or post‐dilation and allows for full repositioning and redeployment. The valve a paravalvular seal that has shown great outcomes in reducing paravalvular leakage [35, 36]. Preliminary results of the RESPOND trial demonstrated good clinical outcomes and the effectiveness of the Lotus Valve in the treatment of BAV patients at 30-days post-implantation [37]. A recent study, demonstrated good clinical outcomes for the Lotus valve in the treatment of BAV patients in terms of valve circularity and hemodynamics, without significant paravalvular regurgitation [35]. However, this clinical study only examined three patients and the authors highlighted the uncertainty in the potential of improved outcomes given the use of a different valve size [35]. TAV sizing for BAV patients is currently based on those used for tricuspid aortic valves using perimeter-derived and area-derived diameters measured from MSCT scans. However, certain patients can fall into the criteria for more than one valve size (Table 1), and so the effects of over- and under-sizing requires further investigation.

**Table 1:** Sizing chart for transcatheter valves [57].

|  | Portico | Lotus | Acurate | Engager |
| --- | --- | --- | --- | --- |
| Valve sizes, mm | 23/25 | 23/25/27 | 23/25/27 | 23/26 |
| Aortic annulus dimension, mm | 19–21 (23) | 20–23 (23) | 20–23 (23) | 21–27 |
|  | 21–23 (25) | 23–25 (25) | 23–25 (25) |  |
|  |  | 25–27 (27) | 25–27 (27) |  |
| Sheath size, Fr | 18 | 18/20 | 18 | 30 |
| Leaflet tissue | Bovine | Bovine | Porcine | Bovine |
| Repositionable | Partial | Complete | Partial | Partial |

In this study, a patient-specific finite element (FE) model of a BAV stenosis who was previously treated using a 25 mm Lotus valve but fits the criteria between a 25 mm and 27 mm Lotus valve, was developed using MSCT images. This model was applied to compare the post-deployment configuration of an undersized or oversized Lotus valve (25 mm and 27 mm respectively) and the biomechanical interaction of each valve size with the aortic root tissue, to provide an advanced understanding and inform sizing criteria for a BAV patient.

## Materials and Methods

### Aortic Root Model

A patient-specific model of a stenosed BAV (Type I) was reconstructed from MSCT images with slice thickness of 0.625 mm, slice dimensions of 512 x 512 and pixel spacing of 0.664 mm. Mimics 14.1 Imaging Software (Materialise, Leuven, Belgium) was used to threshold the leaflets, calcifications, and aortic root. 3 Matic (Materialise, Leuven, Belgium) and TetGen (WIAS, Berlin, Germany) were used to generate volume meshes of the aortic root and calcifications and leaflets. The geometries were then imported into Abaqus Explicit 6.13 (SIMULIA, Providence, RI). An assembly was then generated matching the corresponding nodal positions of the intersecting surfaces.

### Lotus™ Valve Model

Two different models of Lotus valve geometry; 25 mm and 27 mm were created. The braid geometry was imported into Abaqus 6.13 as a wire part and meshed using 3-noded quadratic beam elements (B32). The wire overlap within the braid was modelled using spring connector elements.

### Constitutive Models

The Lotus valve braid was modelled using a superelastic material inbuilt user subroutine (VUMAT), correlated to the experimental crush test of a nitinol Lotus valve braid as described in [38].

The ascending aorta and the aortic sinus were modelled as isotropic hyperelastic materials. They were modelled using the first order Ogden model [189] fitted to uniaxial test data of human tissue [40, 190]. The aortic valve leaflets were modelled using a linear elastic model with Young’s modulus of 1.6 MPa, a Poisson’s ratio of 0.495 and a density of 1140 kg/m^3^ [40, 194]. The calcification was modelled as a linear elastic material with Young’s modulus 10 MPa, Poisson’s ratio 0.35 and a density of 2000 kg/m^3^ [39, 40].

### Boundary and Loading Conditions

The crimping and deployment of the two valve sizes (25 mm and 27 mm) into the BAV geometry was modelled in Abaqus Explicit 6.13 (SIMULIA, Providence, RI). The valves were crimped inward radially using zero friction contact. The valves were then unsheathed into the aortic root by applying an axial displacement to the crimper. During stent deployment, a coefficient of friction of 0.36 was used between the TAV and the calcifications [41], and the interaction between the TAV and the aortic sinus was modelled using a coefficient of friction of 0.1 [42]. A third loading step was applied to the braid to simulate locking of the device using six connector elements displaced inward to ensure the valve was locked to a height of 19 mm.

## Results

The results of the 25 mm model were compared to real-life patient angiograms of the 25 mm BAV patient case. Figure 2 shows the model predictions compared to angiogram images of the valve post-deployment. Measurements of the diameter were taken at 3 planes along the valve, which can be seen in Table 2, show the correlation between the model geometry and the real-life case, with percentage errors averaging <3% in the braid dimensions measured.

**Figure 2:**
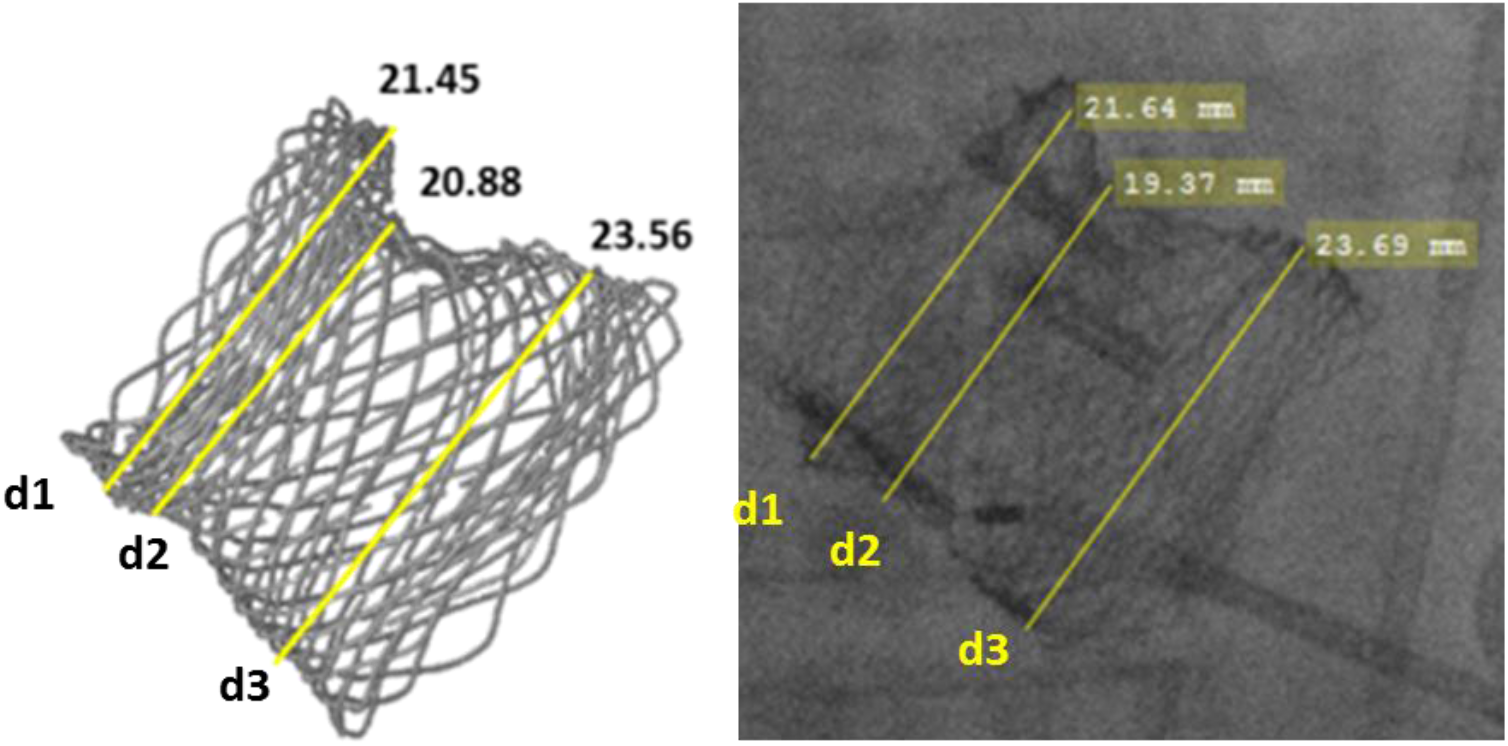
Comparison of model braid geometry with that of post-deployment angiogram from the real-life case. Measurements d1, d2, and d3 are the diameters at plane 1, 2 and 3 respectively.

**Table 2:** Comparison of model valve braid dimensions with that of post-deployment angiogram from the real-life case.

|  | **Model**<br>**Dimensions**<br>**(mm)** | **Angio**<br>**Dimensions**<br>**(mm)** | **Percentage**<br>**Difference (%)** |
| --- | --- | --- | --- |
| **Diameter at**<br>**Plane 1 (d1)** | 21.45 | 21.36 | 0.42 |
| **Diameter at**<br>**Plane 2 (d2)** | 20.88 | 19.37 | 7.23 |
| **Diameter at**<br>**Plane 3 (d3)** | 23.56 | 23.69 | 0.55 |

Figure 2 demonstrates the ability of the model to accurately predict the final deployment geometry of the Lotus valve with an almost identical representation of the reduction of the braid’s waist. The “waist” of the braid can be observed below d2 where the braid reduces in diameter at the mid-section of the braid.

### Valve Eccentricity

Valve eccentricity was firstly examined as elliptical deployment geometries are a complication associated with BAV patients [18] and are associated with leaflet distortion. Leaflet distortion can alter leaflet kinematics and fluid mechanics and ultimately lead to accelerated fatigue of the valve leaflets [43–46]. This is of particular concern with BAV patients due to the limited data existing on long-term durability in BAV cases and the younger patient cohorts associated with BAVs [10]. Eccentricities were measured at the inflow segment of the valve, as this is where the leaflet attachment points reside and represent the highest area of leaflet distortion. The eccentricity of the valve was measured using [47]:

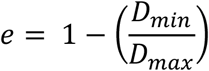

where *D*_*max*_ and *D*_*min*_ are the major and minor axes of the ellipse respectively and zero is the optimal eccentricity.

The valve eccentricities for all models can be found in Table 3. At the basal leaflet attachments, the eccentricity in the 25 mm valve was predicted to be 0.152, whereas an eccentricity of 0.195 was predicted for the 27 mm case.

**Table 3:**
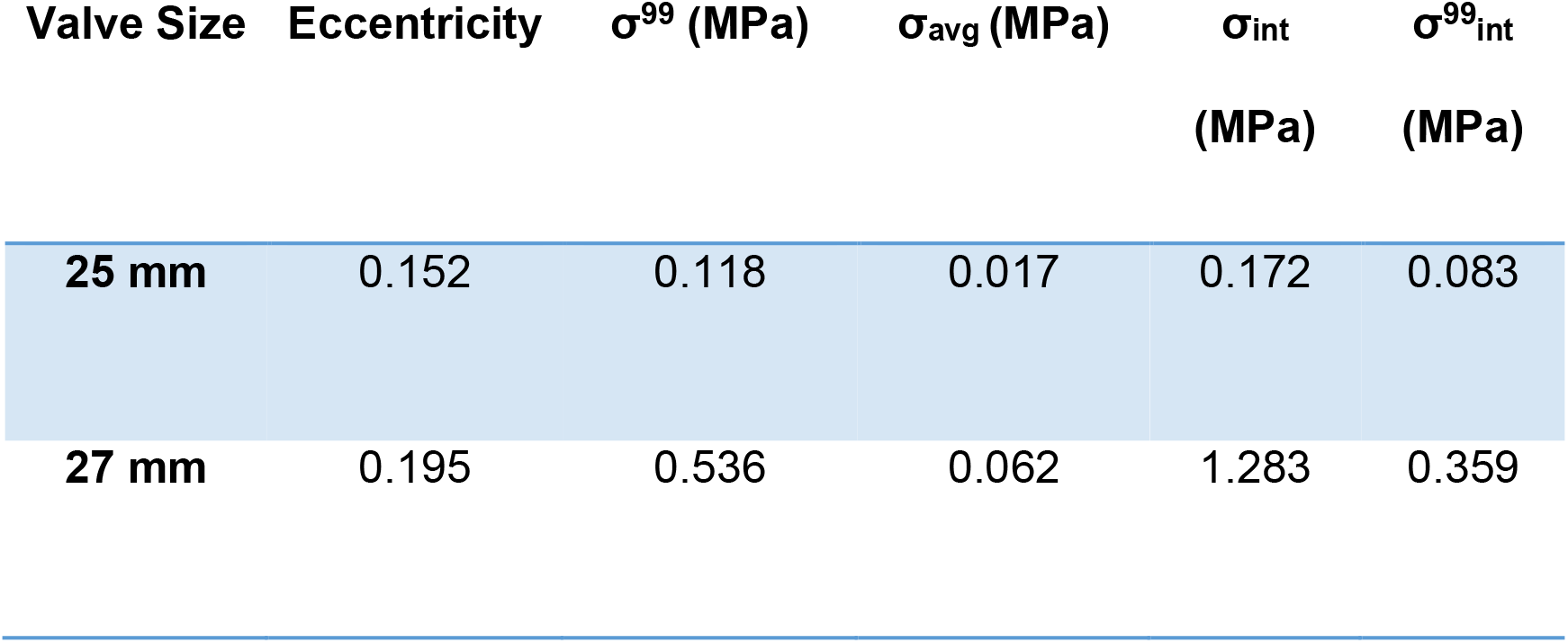
Table of results comparing eccentricity and von Mises stress between the two cases.

### Tissue Stress

The 99^th^ percentile (σ^99^) and the average (σ_avg_) von Mises stress in the aortic sinus were examined for both valve sizes (Table 3). It was predicted that the 27 mm valve had the higher σ^99^ (0.536 MPa) and σ_avg_ (0.118 MPa) when compared to the 25 mm valve case where σ^99^ and σ_avg_ were predicted as 0.118 MPa and 0.017 MPa respectively. Contour plots of the stress in the tissue can be seen in Figure 3 where the 27 mm valve shows higher percentage volume of tissue at higher von Mises stress in comparison to the 25 mm case. The stress in the interleaflet triangle was also examined. The bundle of His is an extension of the atrioventricular node and consists of a branch of specialised cells that facilitate electrical conduction and is positioned at the base of the interleaflet triangle [48–51]. Previous studies have proposed that injury in this area is an underlying cause of conductance interference [49, 50]. Both the maximum and 99^th^ percentile stress in the interleaflet triangle were also compared for both valve sizes with the 25 mm valve showing lower stress (0.172,0.083 MPa) when compared to the 27 mm valve (1.283, 0.359 MPa).

**Figure 3:**
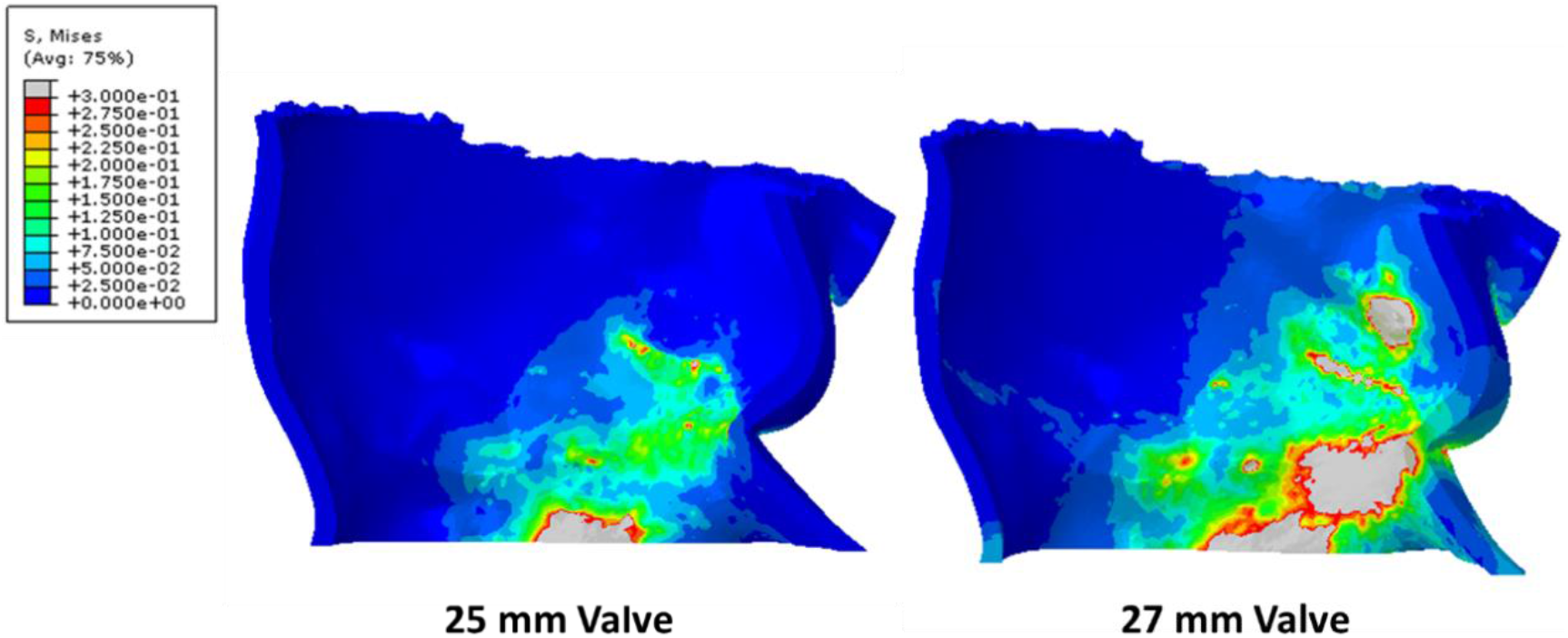
Contour plots von Mises stress in the aortic sinus (MPa) post deployment the 25 mm and 27 mm valve.

### Areas of Mal-Apposition

Although under-sizing of valves reduces stress in the surrounding native aortic root current TAVI guidelines suggest upsizing of the device in order to reduce volumes of PVL [29–31]. As BAV patients are commonly associated with severe PVL [9, 15] “gaps” or mal-apposition of the valve to surrounding tissue which may be potential areas of PVL were examined qualitatively using similar methods to those implemented in [52]. Figure 4 demonstrates the areas of mal-apposition for both valve sizes. There does not appear to be a considerable difference in the regions where mal-apposition occurs reported for both valve cases. It must be noted the Lotus valve includes a paravalvular seal which has been designed to significantly reduce gaps between the stent an tissue [53]. In the absence of modelling the paravalvular seal, mal-apposition can only be compared qualitatively between the two valves.

**Figure 4:**
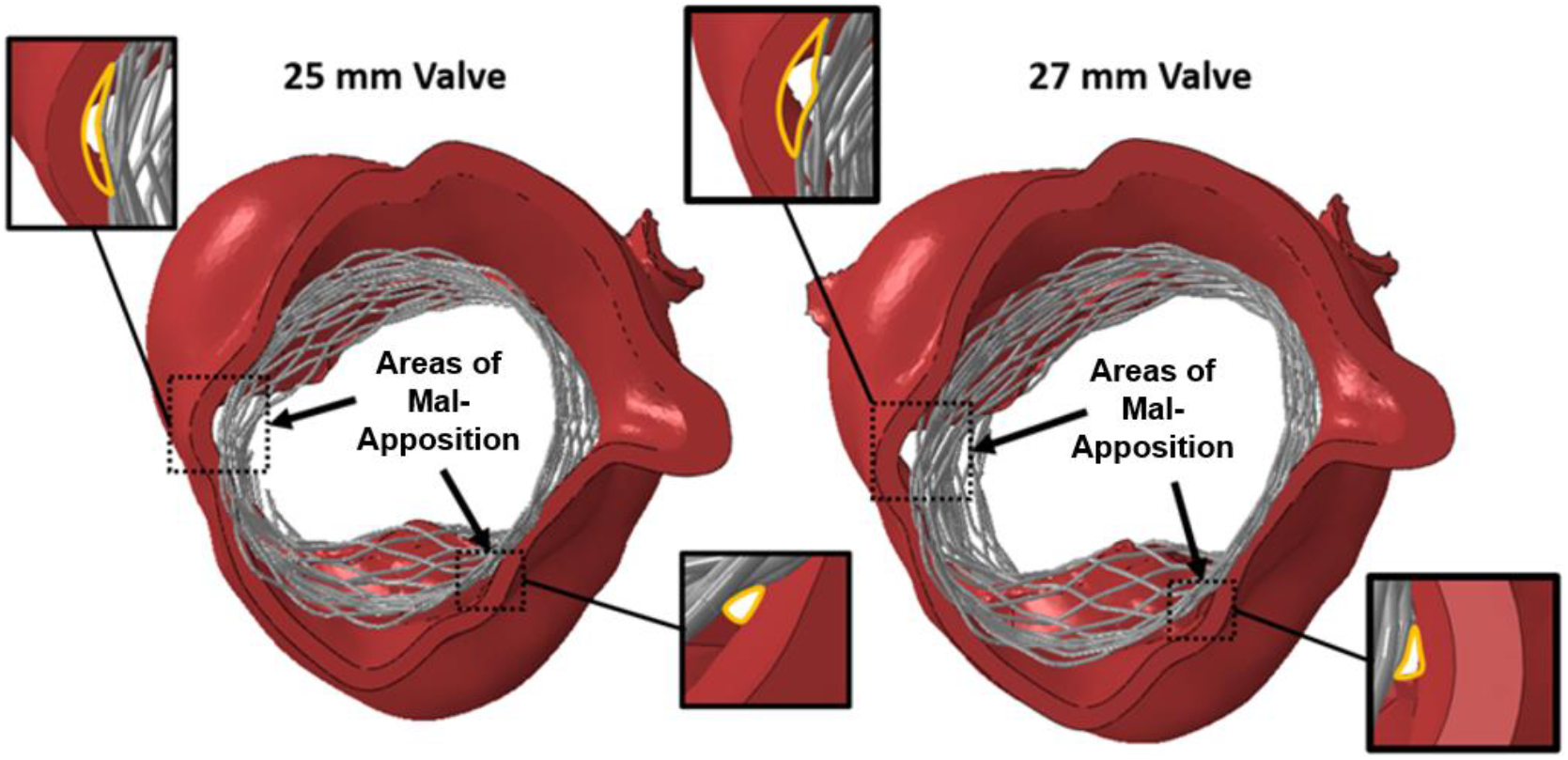
Schematic of the valve braid and aortic root with highlighted areas of mal-apposition.

## Discussion

In this study, patient-specific modelling was used to investigate a patient-specific case of a BAV patient treated with the Lotus valve. The results of this model have shown near perfect correlation to images of the real-life patient case post-TAVI. This model has been used to investigate TAV sizing for BAV patients using the Lotus valve by comparing the two different valve sizes (25 mm and 27 mm) in the same patient anatomy in terms of valve eccentricity and tissue stress. These results suggest that the 25 mm valve was optimal in both lowering eccentricity and tissue stress when compared to the oversized 27 mm valve. The eccentricities reported for the Lotus valve post-deployment in the BAV patient in this study are comparable to those reported in the treatment of tricuspid aortic valve stenosis using other valve types [40, 54].

The limitations to this study must be noted, namely; the uses of isotropic material properties over anisotropic properties, aortic pressure and pre-stress, and that the model examines static only initial deployment conditions. In the absence of modelling the paravalvular seal, an accurate representation of the areas of mal-apposition with potential for PVL cannot be accurately quantified. However, it is reasonable to assume the paravalvular seal would have a similar impact in reducing mal-apposition for both valve sizes. In this study, the assessment of mal-apposition was used solely as a comparative measure between the two valves using similar methods to those previously used by Wang *et al*. in assessing the potential for PVL [52]. Finally, in this study only one patient-specific anatomy was examined, and as such the specific results regarding the relationship between valve sizing, eccentricity and aortic tissue stress cannot be assumed to represent the entire population. However, this is the first detailed FE analysis to provide an understanding of the impact of different TAV sizing in BAV patients.

It was predicted in this study that the braid was more eccentric in the 27 mm case with an eccentricity of 0.195 at the basal plane. This compared to an eccentricity of 0.152 in the 25 mm case. Clinically it has been found that bicuspid patient anatomies have been shown to lead to more eccentric valve geometries [16, 18]. The eccentricity of the native aortic root for the BAV patient in this study is 0.143. The eccentricities observed in this patient are higher than what has previously been reported for the Lotus valve in tricuspid aortic valve patients [47]. The unique mechanism that locks the valve braid leads to desirable eccentricities with average eccentricities of 0.06 ± 0.04 reported clinically for tricuspid aortic valve patients [47]. However, it must be noted that levels of eccentricity of 0.18 and 0.25 have been computationally predicted for the CoreValve and the Edwards SAPIEN Valve in clinical cases of tricuspid aortic valve patients [40, 54]. These results indicate that the locked braid of the Lotus valve is suitable in the treatment of eccentric bicuspid anatomies.

Using a combination of FEA and CFD analysis of a generic TAV geometry it has been predicted that valve eccentricities greater than 0.134 lead to backflow leakage, whereas an eccentricity of 0.267 was predicted to increase the peak leaflet stress by 143% [55]. Although the eccentricity of 0.152 reported for the 25 mm valve and 0.195 reported for the 27 mm valve are acceptable in terms of what has been reported clinically, they exceed the threshold eccentricity of associated with backflow reported by [55]. It must be noted however, that leaflet design is important in valve performance and so the eccentrics reported for a generic TAV geometry [55] can only be used as an indication of sub-optimal eccentricity and cannot be directly compared to the Lotus valve. This should be considered carefully in relation to the treatment of BAV patients, particularly as TAVI move into the treatment of younger patient cohorts. Nevertheless, the 25 mm valve showed preferable eccentricity over the 27 mm valve and comparable eccentricities to those previously reported in clinical cases of tricuspid aortic valve patients treated with TAVs.

The levels of von Mises stress were compared for both valve sizes. The 25 mm case shows lower stress in the sinus when compared to the 27 mm valve case, however, the σ^99^ is below the ultimate tensile stress of the aortic sinus (2.3 to 3.1 MPa) in both cases [56]. The values of maximum and 99^th^ percentile peak stress in the interleaflet triangle were 0.172 MPa and 0.083 MPa for the 25 mm valve and 1.283 MPa and 0.359 MPa for the 27 mm valve. As higher stress in this region have been associated with conductance abnormalities [49, 50] these results suggest lower potential for conductance abnormalities in the 25mm valve case.

The results of this study suggest that the 25 mm valve was more suitable in terms of reduction of stress in the tissue and reduced eccentricity. Current guidelines for TAVI suggest upsizing the device relative to the native annulus to reduce volumes of PVL [29–31]. However, Figure 4 shows cross sections of the valve that indicate that for this patient case oversizing does not appear to considerably reduce the degree of mal-apposition that can indicate potential for PVL. Therefore, on the basis of the results of this study, the 25 mm valve size was deemed most appropriate size in terms of reduced eccentricity and stress reduction in the patient-specific anatomy examined.

## Conclusion

In this study, FE models were developed simulating the deployment of a 25 mm and a 27 mm Lotus valve into the patient-specific aortic root geometry of a clinical BAV case to investigate the efficacy of the Lotus valve in the treatment of BAV stenosis and to examine oversizing in a BAV patient case. The results of this study predicted that the eccentricity for this patient was lower than that previously reported computationally for the other valve types in the treatment of tricuspid aortic valve patients. Furthermore, it was shown the 25 mm valve was the most suitable in terms of eccentricity and stress reduction, and also that this sizing did not increase the likelihood of mal-apposition. On the basis of these results traditional annulus measurement and device sizing may be suitable in the case of the Lotus valve.

## Acknowledgements

The authors acknowledge funding from the Irish Researcher Council Enterprise Partnership Scheme Postgraduate Scholarship 2014 in collaboration with Boston Scientific (EPSPG/2014/120) and acknowledge the Irish Centre for High-End Computing (ICHEC).

